# EWAS-Galaxy: a tools suite for population epigenetics integrated into Galaxy

**DOI:** 10.1101/553784

**Authors:** Katarzyna Murat, Björn Grüning, Paulina Wiktoria Poterlowicz, Gillian Westgate, Desmond J Tobin, Krzysztof Poterlowicz

## Abstract

**Background:** Epigenome-wide association studies (EWAS) analyse genome-wide activity of epigenetic marks in cohorts of different individuals to find associations between epigenetic variation and phenotype. One of the most common technique used in EWAS studies is the Infinium Methylation Assay, which quantifies the DNA methylation level of over 450k loci. Although a number of bioinformatics tools have been developed to analyse the assay they require some programming skills and experience to use them.

**Results:** We have developed a collection of user-friendly tools for the Galaxy platform for those without experience aimed at DNA methylation analysis using the Infinium Methylation Assay. Our tool suite is integrated into Galaxy (http://galaxyproject.org), web based platform. This allows users to analyse data from the Infinium Methylation Assay in the easiest possible way.

**Conclusions:** The EWAS suite provides a group of integrated tools that combine analytical methods into a range of handy analysis pipelines. Our tool suite is available from the Galaxy test toolshed, GitHub repository and also as a Docker image. The aim of this project is to make EWAS analysis more flexible and accessible to everyone.

## Background

Over the last several years comprehensive sequencing data sets have been generated, allowing analysis of genome-wide activity in cohorts of different individuals to be increasingly available. Finding associations between epigenetic variation and phenotype is a significant challenge in biomedical research. Recently performed genome-wide association studies (GWAS) have identified variation naturally occurring in the genome associated with disease risk and prognosis, including tumour pathogenesis [16]. This raised interest in the concept of epigenome-wide association studies (EWAS). Epigenome-wide association studies (EWAS) are the solution to exploring and understanding how interactions between genetic background and the environment could affect human health [8]. The term Epigenome means “on top of” the genome and refers to specific changes in genome regulatory activity occurring in response to environmental stimuli [26]. Epigenetic modifications do not change the underlying DNA sequence, but can cause multiple changes in gene expression and cellular function [8]. Some of the epigenetic modifications such as DNA methylation have been described as related to oncogenesis in a range of cancers including one of the deadliest-melanoma [16]. In humans, DNA methylation occurs by attaching a methyl group to the cytosine residue. This has been suggested as a suppressor of gene expression [14]. Multiple methods for DNA methylation analysis were developed, including the polymerase chain reaction (PCR) and pyrosequencing of bisulfite converted DNA, dedicated to study a small number of methylation sites across a number of samples [27]. Assays like whole genome bisulfite sequencing (WGBS) and reduced representation bisulfite sequencing (RRBS) allow global quantification of DNA methylation levels. However, running this type of analysis for a larger number of samples can be prohibitively laborious and expensive [15]. The Illumina Methylation Assay [12] offers unprecedented applicability and affordability due to the low costs of reagents, short time of processing, high accuracy and low input DNA requirements. It determines quantitative array-based methylation measurements at the single-CpG-site level of over 450k loci [25] covering most of the promoters and also numerous other loci. The makes assay suitable for systematic investigation of methylation changes in normal and diseased cells [26]. As such it has become one of the most comprehensive solutions on the market [17]. However, Illumina Genome Studio is not suitable for everyone and as a commercial software generates additional costs. Therefore there is a need to create freely available software able to perform comprehensive analysis including quality control, normalisation and detection of differential-methylated regions [17]. Open-source software packages (e.g. DMRcate [24], Minfi [10], ChAMP [18], methylumi [7], RnBeads [3]) require high performance computational hardware as well as command line experience in order to run the analysis. This is why one of aims of the our EWAS pipeline was to set and implement these methods into user-friendly environment. An EWAS suite (summarized in table 1) developed to provide users with an enhanced understanding of the Infinium Methylation Assay analysis tool. The tool suite includes methods for preprocessing with stratified quantile normalisation **minfi_ppquantile** or extended implementation of functional normalisation **minfi_ppfun** with unwanted variation removal, sample specific quality assessment **minfi_qc** and methodology for calling differentially-methylated regions and sites **minfi_dmr** and positions detection **minfi_dmp**. All scripts were wrapped into a web based platform - Galaxy, a graphical interface with tools, ready to run workflows providing a solution for non-programmer scientists to analyse their data and share their experience with others [9]. Configuration files are publicly published on our GitHub repository [22] with scripts and dependencies settings also available to download and install via Galaxy test toolshed [21]. Our suite was created and tested using a Planemo workspace with a default configuration and shed tool setup available via Docker (operating-system-level virtualization) [22].

**Table 1.** Summary of the EWAS suite tools inputs and outputs

| Tool ID | Input | Output | Description |
| --- | --- | --- | --- |
| minfi_read450k | IDAT | RGChannelSet | read the .IDAT files |
| minfi_mset | RGChannelSet | MethylSet | convert the Red/Green .IDAT's for an Illumina methylation array |
| minfi_qc | MethylSet /GenomicMethylSet | DataFrame | quality assessment |
| minfi_rset | MethylSet/GenomicRatioSet | RatioSet | converting methylation data from methylation and unmethylation channels, to ratios (Beta and M-values) |
| minfi_ppfun | RGChannelSet | GenomicRatioSet | functional normalization preprocessing |
| minfi_ppquantile | RGChannelSet/GenomicMethylSet | GenomicRatioSet | stratified quantile normalization |
| minfi_maptogenome | MethylSet/RGChannelSet/RatioSet | GenomicRatioSet | add genomic coordinates to each probe together with some additional annotation information |
| minfi_geo | GEO accession | GenomicRatioSet | download data from GEO database |
| minfi_getbeta | MethylSet/RatioSet/GenomicRatioSet | DataFrame | return Beta value |
| minfi_getCN | MethylSet/RatioSet/GenomicRatioSet | DataFrame | return coordinating node |
| minfi_getM | MethylSet/RatioSet/GenomicRatioSet | DataFrame | return the Fisher information corresponding to a model and a design |
| minfi_pheno | RatioSet/GenomicRatioSet | DataFrame | extract phenotype data |
| minfi_getanno | GenomicRatioSet | DataFrame | access provided annotation |
| minfi_getsnp | GenomicRatioSet | DataFrame | return SNP information of the probes |
| minfi_dropsnp | GenomicRatioSet | GenomicRatioSet | drop the probes that contain either a SNP at the metylated loci interrogation or at the single nucleotide extension |
| minfi_dmp | MethylSet/GenomicRatioSet | DataFrame | return differentially-methylated positions |
| minfi_dmr | GenomicRatioSet | DataFrame | return differentially-methylated regions |

## Tools Description

The workflow combines 7 main steps (see Figure 1), starting with raw intensity data loading (.idat) and then preprocessing and optional normalisation of the data. The next quality control step performs an additional sample check to remove low-quality data, which normalisation cannot detect. The workflow gives the user the opportunity to perform any of these preparation and data cleaning steps, including the next highly recommended genetic variation annotation step resulting in single nucleotide polymorphism identification and removal. Finally, the dataset generated through all of these steps can be used to hunt (find) differentially-methylated positions (DMP) and regions (DMR) with respect to a phenotype covariate. Functional annotation of data generates clinically meaningful information about methylation changes with visual representation of these genes and functions. All the tools and single preparation and analysis steps are shown in Figure 2 and explained in detail below.

**Figure 1.**
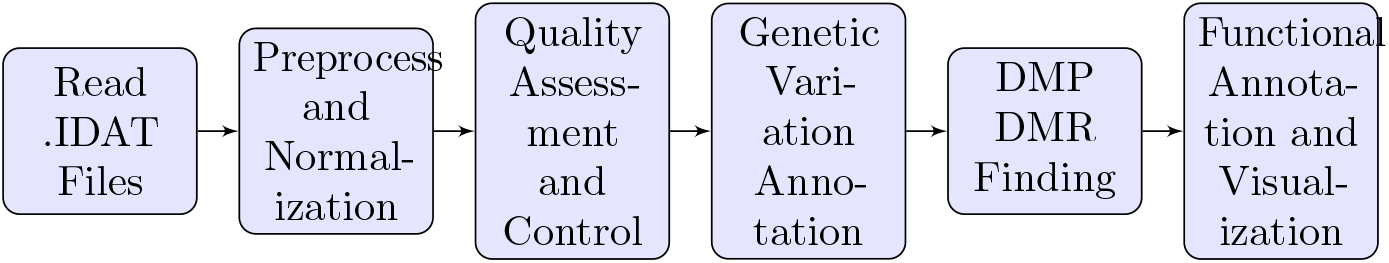
Simplified workflow for analysing epigenetics data

**Figure 2.**
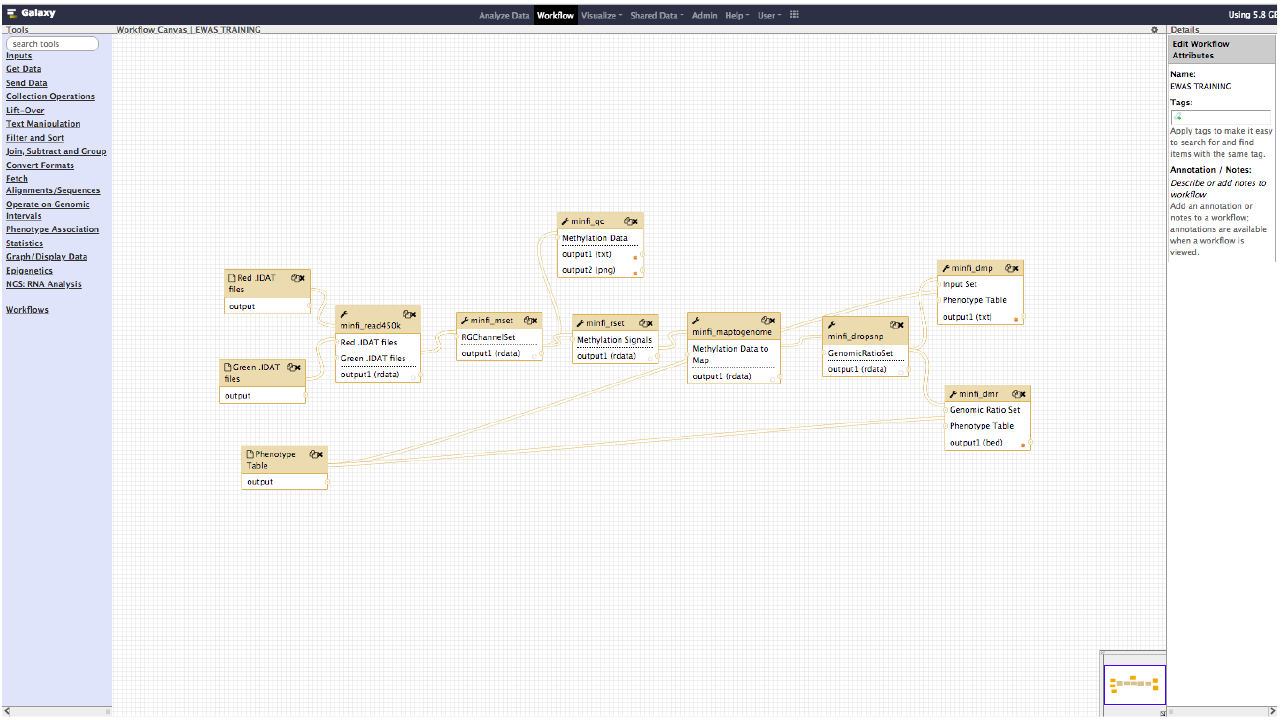
Screenshot from the Galaxy Workflow Editor, showing EWAS example workflow as discussed in the Analyses section.

### Data Loading

The 450k assay interrogates fluorescent signals (qreen and red) from the methylated and unmethylated sites into binary values which can be read directly as IDAT files [12]. Illumina’s GenomeStudio solution converts the data into plain-text ASCII files losing a large amount of information during this process [1]. To prevent this kind of data loss we developed an R based tool **minfi_read450k** which is a combination of illuminaio readIDAT and minfi RGChannelSet functions. The tool loads intensity information from both treatment and control data and based on this it builds up a RGChannelSet class.

### Preprocessing and Normalization

RGChannelSet represents two colour data with a green and a red channel and can be converted into methylated and unmethylated signals assigned to MethylSet or Beta values. Betas build in RatioSet object, and estimate the methylation level using channels ratio in a range between 0 and 1 with 0 being unmethylated and 1 being fully methylated [1]. Users can convert from RGChannelSet into a MethylSet using the **minfi_mset** tool or compute Beta values using **minfi_rset** tool, if no normalisation is performed. However, these two classes can also be preprocessed and normalised with two methods avaliable [1]. **Minfi_ppquantile** implements stratified quantile normalisation preprocessing and is supported for small changes like in one-type samples e.g. blood datasets. In contrast, **minfi_ppfun** is aimed at global biological differences such as healthy and occurred datasets with different tissue and cell types. This is called the between-array normalisation method and removes unwanted variation [1]. Both of these methods return GenomicRatioSet class, that holds comprehensive information about methylation assays mapped to a genomic location [1].

### Quality Assessment and Control

Data quality assurance is an important step in Infinium Methylation Assay analysis. The **minfi_qc** tool extracts and plots the quality control data frame with two columns mMed and uMed which are the medians of MethylSet signals (Meth and Unmeth). Comparing these against one another allows users to detect and remove low-quality samples that normalisation cannot correct [10].

### Annotating probes affected by genetic variation

Single nucleotide polymorphism (SNP) regions may affect results of downstream analysis. **Minfi_getsnp** return data frames containing the SNP information of unwanted probes to be removed by **minfi_dropsnp** tool [1].

### DMPs and DMRs Identification

The main goal of the EWAS suite is to simplify the way differentially-methylated loci sites are detected. The EWAS suite contains a **minfi_dmp** tool detecting differentially-methylated positions (DMPs) with respect to the phenotype covariate, and **minfi_dmr** provides a solution for finding differentially-methylated region (DMRs) [10]. DMRs can be tracked using a bump hunting algorithm. The algorithm first implements a t-statistic at each methylated loci location, with optional smoothing, then groups probe into clusters with a maximum location gap and a cutoff size to refer the lowest possible value of genomic profile hunted by our tool [13].

### Functional Annotation and Visualization

In addition to downstream analysis, users can access annotations provided via Illumina (**minfi_getanno**) [1] or perform additional functional annotations using the Gene Ontology (GO) tool (**clusterprofiler_go**). The Gene Ontology (GO) tools provides a very detailed representation of functional relationships between biological processes, molecular function and cellular components across data [6]. Once specific regions have been chosen, **clusterprofiler_go** visualize enrichment result (see Figure 5). Many researchers use annotation analysis to characterise the function of genes, which highlights the potential for Galaxy to be a solution for wide-ranging multi-omics research.

### Documentation and Training

We have also provided training sessions and interactive tours for user self-learning. The training materials are freely accessible at the Galaxy project Github repository [19]. Such training and tours guide users through an entire analysis. The following steps and notes help users to explore and better understand the concept. Slides and hands-on instruction describes the analysis workflow, all necessary input files are ready-to-use via Zenodo [20], as well as a Galaxy Interactive Tour, and a tailor-made Galaxy Docker image for the corresponding data analysis.

## Potential implications

Increased interest in skin cancer biomarker identification led us to validate the differentially-methylated regions analysis using the Illumina 450K Methylation array data of melanoma biopsies pre and post MAPKi treatment [11], obtained from the Gene Expression Omnibus (GEO) (GSE65183). Methylation profiling by genome tiling array in melanoma can help us understand how non-genomic and immune changes can have an impact on treatment efficiency and disease progression. Raw image IDAT files were loaded into the Galaxy environment using Data Libraries. EWAS workflow was run on Red and Green dataset collections of patient-matched melanoma tumours biopsied before therapy and during disease progression. The IDAT files, pre-defined phenotype tables and up-to-date genome tables (UCSC Main on Human hg19 Methyl450) [22] were used as inputs. In order to detect poorly performing samples we ran quality diagnostics with **minfi_qc tool**. The provided samples passed the quality control test (on figure 3) as they clustered together with higher median intensities confirming their good quality [1].

**Figure 3.**
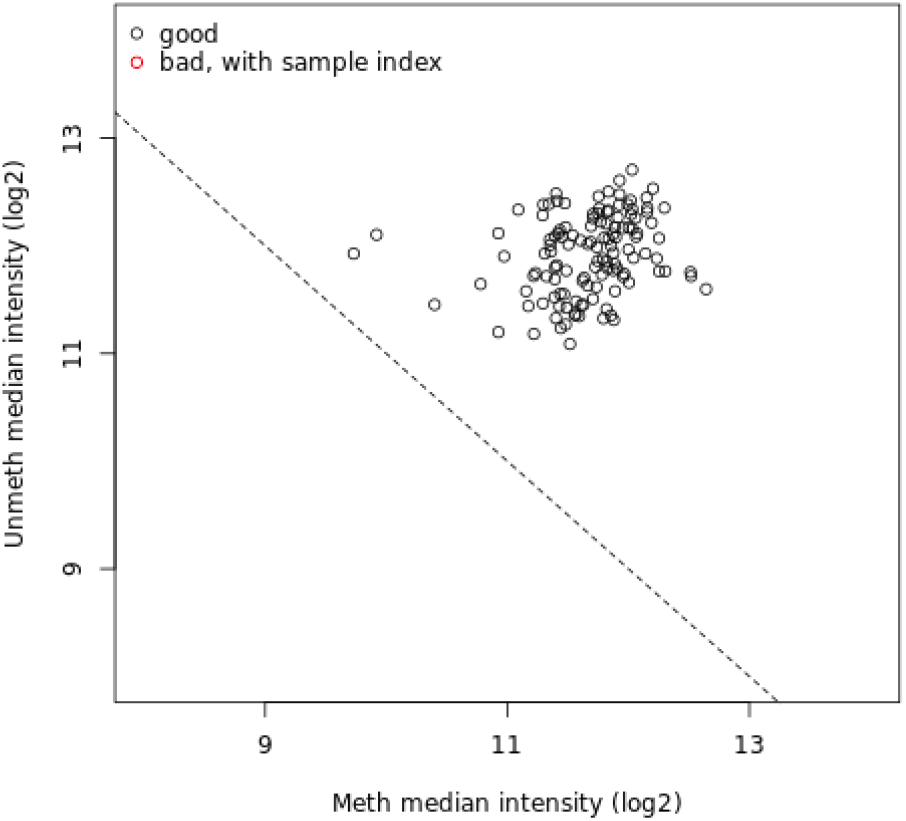
Quality Control Plot representation of melanoma pre and post MAPKi treatment samples.

Differentially-methylated loci were identified using single probe analysis implemented by **minf_dmp** tool with the following parameters: phenotype set as **categorical** and qCutoff size set to **1**. The bump hunting algorithm was applied into the **minfi_dmr** tool to identify differentially-methylated regions (DMRs) with maximum location gap parameter set to **250**, genomic profile above the cutoff equal to **0.1**, number of resamples set to **0**, null method set to **permutation** and verbose equal **FALSE** which means that no additional progress information will be printed. Differentially-Methylated Regions and Positions revealed the need for further investigation of tissue diversity in response to environmental changes [4]. Nearest transcription start sites (TSS) and enhancer elements annotations founded in the gene set can be listed as follows: PITX1, SFRP2, MSX1, MIR21, AXIN2, GREM1, WT1, CBX2, HCK, GTSE1, SNCG, PDPN, PDGFRA, NAF1, FGF5, FOXE1, THBS1, DLK1 and HOX gene family. Although hyper-methylated genes identified by ‘EWAS-suite’ have been previously associated with cancer, this is the first time a link between them and MAPKi treatment resistance is reported. These data demonstrates that PDGFR, which is suggested to be responsible for RAS/MAPK pathway signaling trough activation may regulate the MAPKi mechanism in non responsive tumours. The methylation regulation of this altered status of PDGFR requires additional studies [11]. The PITX1 suppressor gene was found as one of the factors decreasing gene expression in human cutaneous malignant melanoma and might contribute to progression and resistance via promoting cell proliferative activity [23]. It has been found that homeodomain transcription factor MSX1 and CBX2 polycomb protein are likely to be treatment resistance factors and are reported as downregulated and inactivated in melanoma tumours [5]. Previous published studies are limited to local surveys and serial biopsies. Thus, the stimulus of innate or acquired MAPKi resistance may converge on epigenetics.

**Figure 4.**
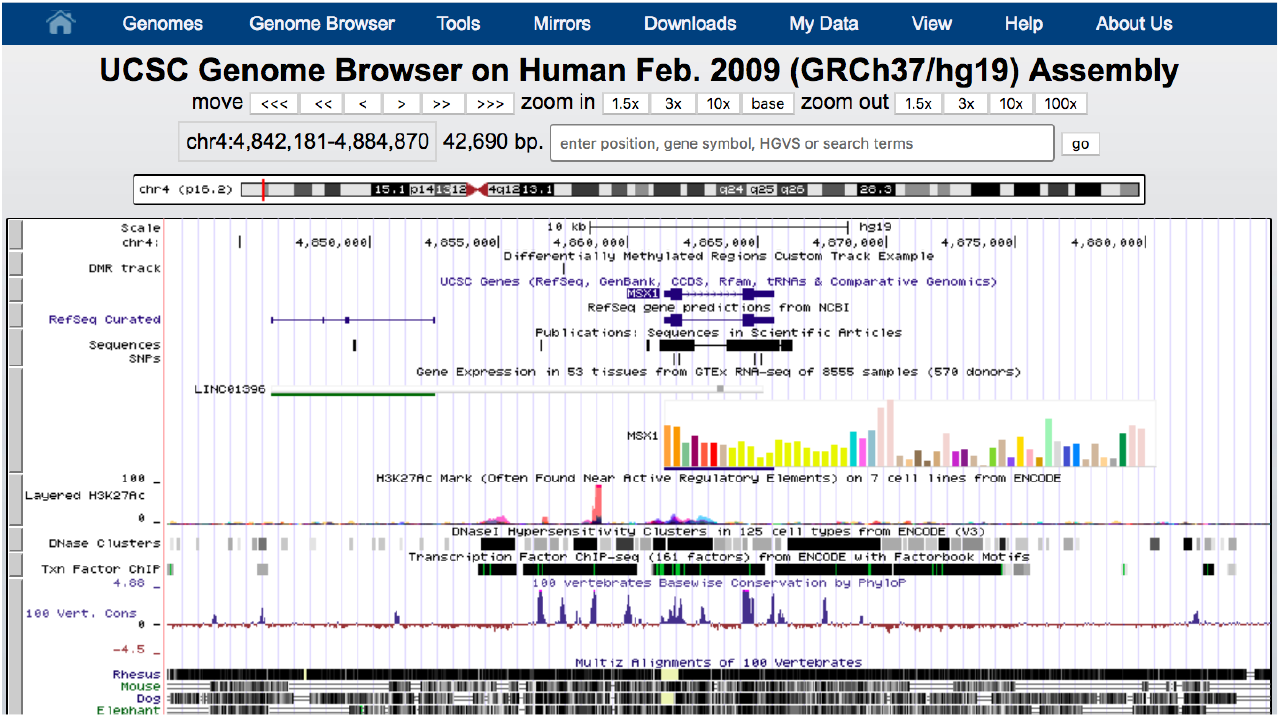
UCSC Example Track.

Functional annotation with GO is a scheme to understand how the annotations are assigned to the genes [2]. These are enrichment GO categories after controlling for false discovery rate (FDR) control figure (see in 5). The greatest significance to the gene output was the pattern specification process (GO:0007389), skeletal system development (GO:0001501) and regionalisation (GO:0003002) meaning that melanoma MAPKi resistance could be related to the cells developmental process within specific environments.

**Figure 5.**
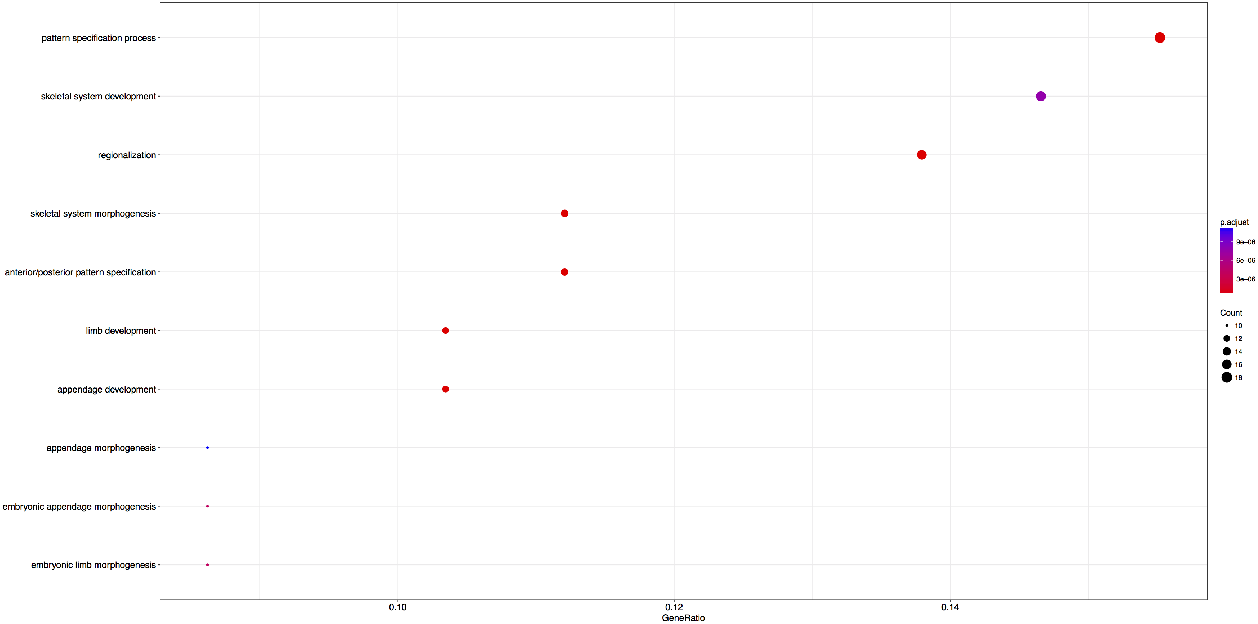
Functional Annotation of DMR’s found in melanoma biopsies pre and post MAPKi treatment.

## Conclusion

With the rapidly increasing volume of epigenetics data available, computer-based analysis of heritable changes in gene expression becomes more and more feasible. Many genome-wide epigenetics studies have focused on generation of the data, however data interpretation is a challenge now. Risk evaluation, disease management and novel therapeutic development are prompting researchers to find novel bioinformatic frameworks and approaches. In this regard we propose a user friendly tool suite available via Galaxy platform ‘EWAS-Galaxy’ This tools suite allows life scientist to run complex epigenetics analyse. [22]. The use case presented provides a tangible example how the EWAS tool suite can provide additional insights into melanoma therapeutic resistance.

## Availability of source code and requirements

Project name: EWAS-Galaxy: a tools suite for epigenomics data analysis integrated into Galaxy

Project home page:https://github.com/kpbioteam/ewas_galaxy
Operating system(s): Linux (recommended), Mac
Programming language: R programming language (version 3.3.2, x86 64bit)
Other requirements: Galaxy [19]
License: License version x

## Availability of supporting data and materials

Test data-set from this article are available in the GEO database under accession GSE65186.

